# Energy-based Modelling of Single Actin Filament Polymerisation Using Bond Graphs

**DOI:** 10.1101/2024.06.14.598965

**Authors:** Peter J. Gawthrop, Michael Pan, Vijay Rajagopal

## Abstract

Bond graphs provide an energy-based methodology for modelling complex systems in a hierarchical fashion; at the moment, the method allows biological system with both chemical and electrical subsystems to be modelled. Herein, the bond graph approach is extended to include chemomechanical transduction thus extending the range of biological systems to be modelled.

Actin filament polymerisation and force generation is used as an example of chemo-mechanical transduction and it is shown that the **TF** (transformer) bond graph component provides a practical, and conceptually simple, alternative to the Brownian ratchet approach of Peskin, Odell, Oster & Mogilner. Furthermore it is shown that the bond graph approach leads to the same equation as the Brownian ratchet approach in the simplest case.

The approach is illustrated by showing that flexibility and non-normal incidence can be modelled by simply adding additional bond graph components and that compliance leads to non-convexity of the force-velocity curve.

Energy flows are fundamental to life; for this reason, the energy based approach is utilised to investigate the power transmission by the actin filament and its corresponding efficiency.

The bond graph model is fitted to experimental data by adjusting model physical parameters.

## 1 Introduction

“Systems Biology explores how parts of biological entities function and interact to give rise to the behaviour of the system as a whole.” [1]. This Systems Biology approach is being used as the conceptual basis of understanding living entities as a system with hierarchical interacting parts, including the molecular, organelle, cellular and organ levels, within the Physiome Project [2, 3]. This endeavour requires a conceptual framework which enables models of parts to be readily integrated to model systems in a modular fashion. The concept of *energy* provides a way of integrating parts in different physical domains such as chemical, electrical, protonic and mechanical; for this reason, the energy-based *Bond Graph* approach has been suggested as a conceptual framework [3, 4].

Bond graphs were introduced (in the context of modelling engineering systems) by Paynter [5, 6] to model *energy* transduction within and between different physical domains including electrical, mechanical and hydraulic; a number of introductory texts are available [7–9]. Bond graph models of chemical reaction networks were introduced in the seminal papers of Oster et al. [10, 11], and later extended within the context of Systems Biology [12–18]. Bond graph models connecting such chemical reaction networks to the electrical domain within the Systems Biology context have been developed [19–21]; in particular, redox reactions have been modelled [22]. A recent introduction and overview of the bond graph approach to systems biology is available [23] and a short introduction to bond graph modelling is given in the Supplementary Material.

Bond graph models coupling the chemical and electrical domains have already been established [19, 21] and have been used to build systems describing nerve impulse propagation [19], membrane transporters [20], redox reactions and mitochondrial chemiosmosis [22]. Living systems also make use of the coupling between chemical and *mechanical* domains in, for example, muscles [24], bacterial flagellar motors [25] and the molecular motors of the cytoskeleton [26] which also provide the motivation for synthetic molecular motors [27]. However, using the Bond Graph approach to connect the chemical and mechanical domains within the Systems Biology context is an outstanding issue; the purpose of this paper is to resolve this issue using the modelling of an Actin Filament as a particular example of chemomechanical transduction. As shown in the sequel, a contribution of this paper is to shown that chemomechanical transduction has a similar bond graph representation to that of chemoelectrical transduction. This means that the bond graph approach can be used to build energy-based models of biological systems with mechanical components as well as the preexisting chemical and electrical components.

Actin filament mechanics and dynamics govern cell and tissue physiology throughout life. Actin dynamics and organisation play an integral role in the formation of tissues at cell-cell adhesions [28], play a central role in cell division and growth [29], contribute to cell stiffness [30] and interact with myosin to generate contractile forces in motile and muscle cells [31].

The growth and breakdown of actin filaments underlies actin dynamics. The rate of actin polymerisation is tightly regulated by an interconnected network of chemical reactions, and mechanical forces. In a seminal paper on actin-based cellular motility, Peskin et al. [32] introduce the concept of a *Brownian ratchet* to “present a mechanical picture of how polymerising filaments can exert mechanical forces”. The mechanism provides a physical and thermodynamically consistent explanation of how the Gibbs free energy of actin polymerisation is converted into a mechanical pushing force at the leading edge. The Brownian ratchet approach has been embellished in a number of ways to provide a better fit to experimental data and remains at the core of current understanding of how actin polymerisation exert mechanical forces [33–35]. An elegant summary of the work of Peskin, Odell, Oster & Mogilner is given by Pollard [36] who highlights three key papers: [32] which introduces the concept of the Brownian ratchet in the context of a rigid actin filament touching a membrane at a right-angle to the membrane surface; [37] which extends the theory to flexible filaments touching the membrane at an arbitrary angle; and [38] which further extends the theory to include tethered filaments and capped filaments.

The concept of *energy transduction* as a basis for modelling the interface between chemistry and mechanics has a long history [39, 40]. In this study, we present a novel formulation of single actin filament polymerisation within the bond graph modelling framework.

Although our original intention was to derive a bond graph model of the Brownian ratchet, it has proved possible to to use the same simple technique that was used to model chemoelectrical transduction [19, 21] to model chemomechanical transduction. Despite this simplified approach, we show that the bond graph approach produces a force-velocity relationship identical in form to that of the Brownian ratchet although the bond graph model does not assume a Brownian ratchet mechanism. Furthermore, we illustrate how the basic transduction model can be used as a component to model actin filaments that are flexible and not normal to the impinging surface by adding appropriate bond graph components.

The *transformer* **TF** is the bond graph component which provides the link between different physical domains; in the context of systems biology, the **TF** component has been used to link the chemical and electrical domains [19–22] and to represent stoichiometry [23]. This paper suggests that the **TF** component can be utilised in place of the Brownian ratchet as the link between the chemical and mechanical domains. In particular, the chemomechanical behaviour of a single actin filament is analysed using the bond graph **TF** approach and the corresponding force-velocity curve is derived for rigid and flexible filaments. It is further shown that compliance leads to non-convex curvature of the force-velocity curve – a phenomenon that has been previously reported in the multi-filament case [41–44]. As discussed in § 6,this has implications for the sensitivity of velocity to force.

§ 2 considers the rigid case normal to the surface and the result is explicitly compared with the Brownian ratchet results of Peskin et al. [32]. § 3 considers the power transfer and efficiency of the energy transduction. § 4 extends the bond graph model to the non-normal case and § 5 further extends the model to the flexible case. § 6 examines how the convexity properties of the actin filament force-velocity curve vary with the compliance of the filament. Comparison with the experimental data of Li et al. [35] (corresponding to that of Bieling et al. [34]) is presented in § 7. A short introduction to bond graph modelling is given in the Supplementary Material.

The Python/Jupyter notebook used to generate the Figures is available at: https://github.com/gawthrop/ActinRod24.

## 2 Bond Graph model of actin chemomechanical transduction

In their seminal paper, Peskin et al. [32] consider a actin filament growing by polymerisation against a membrane. The basic parameters are: a polymerisation rate of *α* monomers s^−1^, a depolymerisation rate of *β* monomers s^−1^, a monomer length of *δ* m and a force acting on the barrier of *F* N. Using a Brownian ratchet approach, Peskin et al. [32] (equation 3) show that the tip velocity *V* m s^−1^ is given by

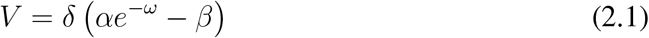

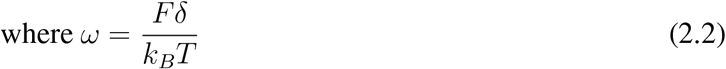

*k*_*B*_ is the Boltzmann constant and *T* the absolute temperature. As indicated in Figure 1(a), there are two physical domains involved: the chemical domain represented by the conceptual chemical reaction *A* ↔ *B* and the mechanical domain represented by a growing actin filament. The left and right sides of the conceptual reaction can be regarded as chemical complexes^1^ where complex

**Figure 1:**
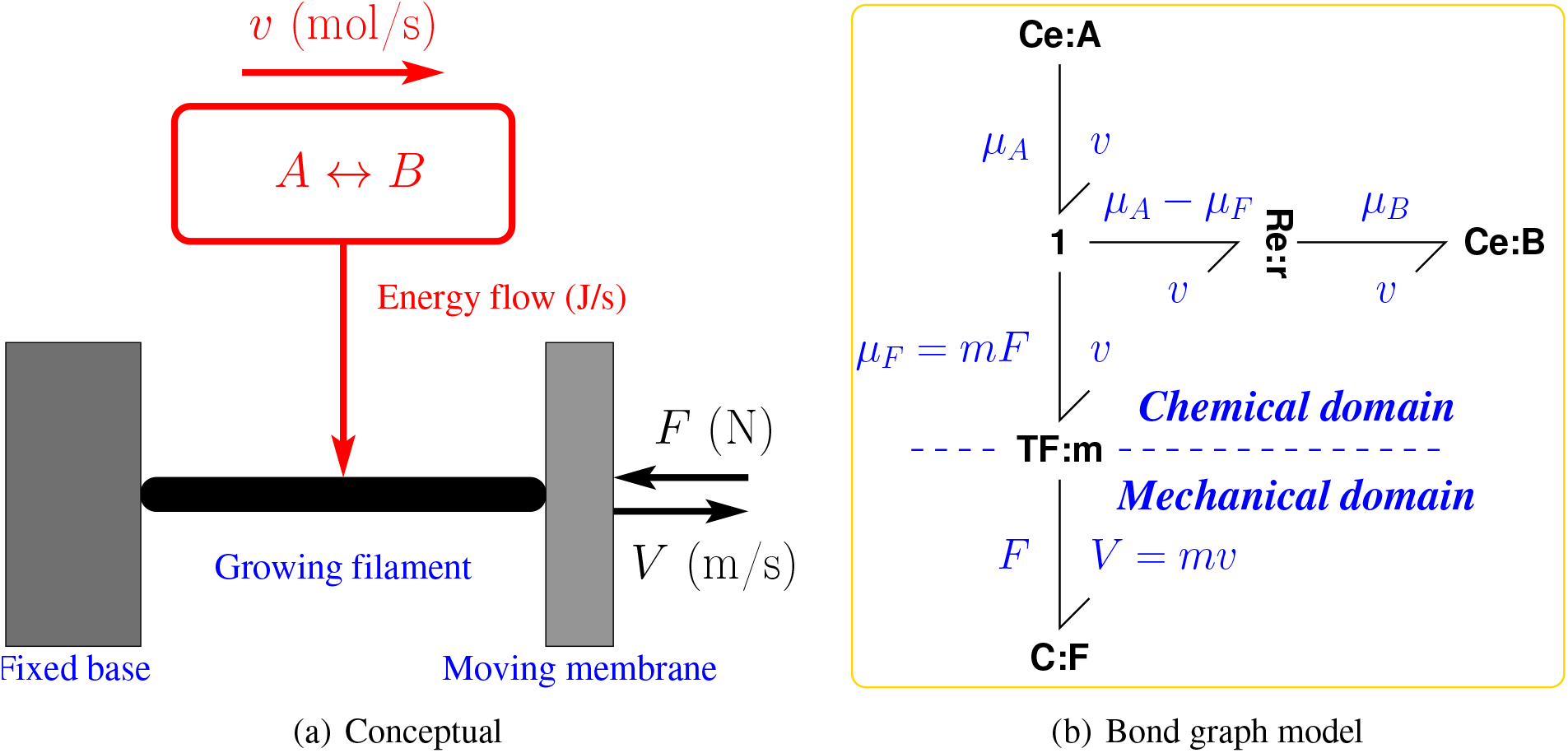
Bond Graph model – rigid filament normal to surface. (a) The chemical reaction A ⇆ B describes the biochemistry of the growing actin filament. The reaction rate *v* (mol s^−1^) also provides the energy flow (J s^−1^) from the chemical to the mechanical domain – chemomechanical transduction. In this section, the actin filament is normal to the membrane. The velocity of the actin tip touching the membrane *V* (m s^−1^) depends on the length and number of monomers added to the polymer per second and is proportional to reaction rate *v*: *V* = *mv*. The constant *m* has units of m mol^−1^ and is derived in § 2.1. The growth of the filament is resisted by a force *F* (N). (b) The bond graph is a precise representation of the energy flows implied by the conceptual diagram where ⇁ corresponds to mechanical or chemical energy flow (J s^−1^). The bond graph **TF**:**m** component represents the chemomechanical transduction with modulus *m* (m mol^−1^). Components **Ce**:**A, Ce**:**B** represent *chemical complexes* (mixtures of species) [15, 45] including ATP hydrolysis; **Ce**:**A** includes the Actin polymer with *n* units and the ATP-bound G-actin monomer; **Ce**:**B** includes the Actin polymer with *n* + 1 units. The bond graph component **Re**:**r** represents the chemical reaction. The *mechanical* spring **C**:**F** represents the force *F* (N) resisting the polymerisation of Actin and the corresponding tip velocity *V* (m s^−1^); this can be thought of as the mechanical equivalent of a chemical chemostat. The annotation in blue is purely explanatory and is not part of the bond graph.

A includes the Actin polymer with *n* units and the ATP-bound G-actin monomer; complex B includes the Actin polymer with *n* + 1 units. As is shown in § 2.2, complexes A and B are related to the *α* and *β* parameters of (2.1). The growth of the actin filament against the membrane can be viewed as *chemomechanical transduction* whereby chemical energy extracted from a conceptual chemical reaction is used to drive the extension of the filament with velocity *V* against the membrane force *F*.

This paper uses a bond graph approach to represent both physical domains together with the chemomechanical transduction. As discussed in the Introduction, the motivation is to be able to include actin dynamics as one component of a modular, energy-based model of a system containing actin filaments. Furthermore, the bond graph approach of this paper yields a simple derivation of (2.1). This section introduces the bond graph material necessary to model the system of Figure 1(a); a more detailed introduction to the bond graph method appears in the Supplementary Material.

Bond graphs unify the modelling of energy within and across multiple physical domains using the concepts of *effort* (*e*) and *flow* (*f*) variables whose product is *power*. In the mechanical domain, the effort variable is force *F* with units of N and the flow variable is velocity *V* with units of m s^−1^ and they have the product *P* = *FV* with units of power J s^−1^. In the chemical domain, the effort variable is *chemical potential µ* with units of J*/*mol and the flow variable is reaction flow rate *v* with units of mol*/*sec; *P* = *µv* also has units of J s^−1^. Bond graphs represent the flow of energy (in any physical domain) between components by the bond symbol ⇁ where the direction of the harpoon corresponds to the direction of positive energy flow. The bond carries the domain-appropriate effort and flow variables; Figure 1(b) has five bonds: four in the chemical domain and one in the mechanical domain. Each bond has been annotated (blue) to show the associated effort and flow variables.

The bond graph **C** component *stores*, but does not dissipate, energy. In the mechanical domain, it corresponds to a spring and the component **C**:**F** is associated the the force *F*. In the chemical domain it corresponds to a chemical species and, because of its special logarithmic constitutive relation, is given the special name **Ce**. In particular **Ce**:**A** is associated with the chemical potential *µ*_*A*_ and **Ce**:**B** is associated with the chemical potential *µ*_*B*_. (As discussed above, A and B are chemical complexes where complex A includes the Actin polymer with *n* units and the ATP-bound G-actin monomer; complex B includes the Actin polymer with *n* + 1 units).

The bond graph **R** component *dissipates*, but does not store, energy. In the mechanical domain, it corresponds to a damper; in the chemical domain it corresponds to a chemical reaction and, because of its special exponential constitutive relation, is given the special name **Re**. The upper part of Figure 1(b) corresponds to the chemical reaction A ⇆ B modulated by the equivalent chemical potential *µ*_*F*_. For the purposes of obtaining the velocity (*V*)–force(*F*) curve, the two **Ce** components are *chemostats* [13, 23] which have fixed values of potential *µ*_*A*_ and *µ*_*B*_.

Electrical components have series (common current) and parallel (common voltage) connections; similarly, the bond graph has two connections represented by the **1** (common flow) and **0** (common effort) junctions. Neither junction dissipates or stores energy; thus, in the case of the **1** junction (with common flow), the efforts sum to zero. In the particular case of Figure 1(b), this implies that the effort on the horizontal bond is the difference between the efforts on the incoming (*µ*_*A*_) and outgoing (*µ*_*B*_) vertical bonds. The bond graph **TF** component is the key to the transformation between physical domains in general and the chemical and mechanical domains in particular; like the junctions, energy is neither dissipated or stored so the power is the same at each of the two connected bonds. This is the indicated in Figure 1(b) where *µ*_*F*_ *v* = *FV*. In particular, the mechanical force *F* is transduced to the equivalent chemical potential *µ*_*F*_ = *mF*. The derivation of the *modulus m* is considered in § 2.1.

It is shown in § 2.2 that the bond graph approach yields an identical force/velocity equation to (2.1) as derived by the Brownian ratchet approach of Peskin et al. [32].

### 2.1 The TF component and modulus

The **TF** component^2^ provides the interface between two different physical domains; in the context of systems biology, the **TF** component has been used to link the chemical and electrical domains [19–22]. The **TF** component is used here to link the chemical and mechanical domains. Although the chemical and mechanical domains differ and correspond to different physical units, each has the same notion of energy measured in Joules (J). The bond graph approach assigns two key variables to each domain: an effort *e* and a flow *f* which have the property that their product is power *p* = *ef* J s^−1^. The chemical domain has chemical potential *µ* J mol^−1^ as effort and molar flow *v* mol s^−1^ as flow and the mechanical domain has force *F* N as effort and velocity *V* m s^−1^ as flow.

Because the **TF** component does not store or dissipate energy, it follows that both effort and flow are related by the same modulus *m*:

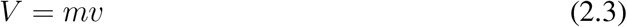

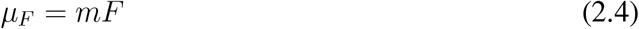

where *µ*_*F*_ is the chemical potential corresponding to the force *F*.

Equation (2.3) converts molar flow into mechanical velocity. As a mole of actin subunits contains *N*_*A*_ molecules where *N*_*A*_ (mol^−1^) is the Avogadro Constant, and assuming that the length of the Actin subunit is *δ* (m), it follows that the **TF** modulus *m* (m mol^−1^) is given by

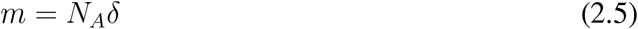

Because *m* appears in both equations (2.3) and (2.4) it follows that the modulus *m* derived from considering the flows can also be used to relate the chemical and mechanical efforts *µ*_*F*_ and *F*.

The use of the **TF** component for chemomechanical transduction is similar to its use for chemoelectrical transduction [19, 21]. Whereas in the chemomechanical case, the modulus *m* transforms mol s^−1^ to m s^−1^, in the chemoelectrical case the modulus *m* transforms mol s^−1^ to C s^−1^ (or, equivalently, A) and *m* is the Faraday constant 9.64 × 10^4^ C mol^−1^ multiplied by the number of electrical charges per molecule.

### 2.2 Comparison with the Brownian ratchet approach

From Figure 1, the net chemical potentials acting on the reaction component **Re**:**r** are *µ*_*A*_ − *µ*_*F*_ and *µ*_*B*_. Hence, the net reaction flow *v* is given the formula [12, 23]:

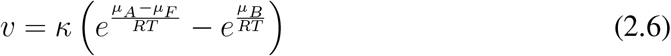

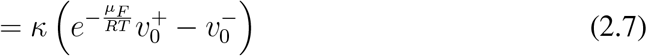

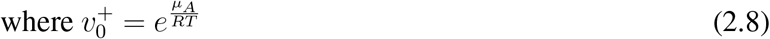

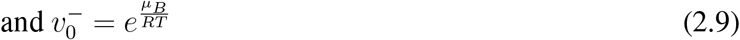

*κ* is the reaction rate constant associated with the reaction component **Re**:**r**, *µ*_*A*_ and *µ*_*B*_ are the chemical potential associated with complexes A and B. *RT* (J mol^−1^) is the product of the universal gas constant *R* (J K^−1^ mol^−1^) and the absolute temperature *T* (K).

Using equations (2.3) – (2.5), the mechanical velocity is:

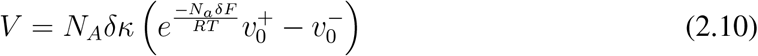

The Bond Graph derived equations (2.10) and the Brownian ratchet equation of Peskin et al. [32, (3)] (2.1) are identical if the following substitutions are made:

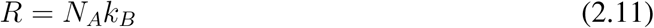

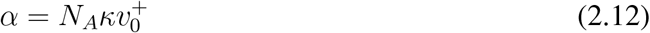

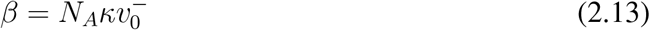

Equation (2.11) is the standard formula for the Gas Constant *R*; Equation (2.12) represents the polymerisation rate in monomers s^−1^ and Equation (2.13) represents the depolymerisation rate in monomers s^−1^.

Thus the formula (2.10) obtained by bond graph arguments and the formula (2.1) obtained by statistical mechanics arguments by Peskin et al. [32] are identical.

### 2.3 Force-velocity curve

Using the *α* − *β* notation, Equation (2.10) becomes:

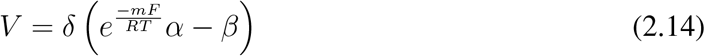

Setting *F* = 0 in equation (2.14), the velocity *V*_0_ corresponding to no membrane force is given by:

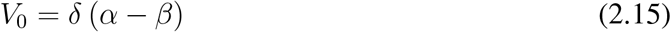

Using the values from Table 1:

**Table 1:** Numerical values from the literature

| Name | Sm. | Value | Unit | Reference |
| --- | --- | --- | --- | --- |
| Polymerisation rate | $\alpha$ | 113 | $\text{s}^{-1}$ | Peskin et al. [32], Fig. 1 |
| Depolymerisation rate | $\beta$ | 1.6 | $\text{s}^{-1}$ | Peskin et al. [32], Fig. 1 |
| Monomer length | $\delta$ | 2.7 | nm | Peskin et al. [32], Fig. 1 |
| Polymer length | $L$ | 30 | nm | Mogilner and Oster [37], Fig. 2 |
| Persistence length | $\lambda$ | 1 | $\mu\text{m}$ | Mogilner and Oster [37], Tab. 1 |
| Contact angle | $\theta_0$ | 90-54=36 | degree | [35], equation (8) |

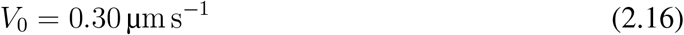

This value is similar to *V*_0_ = 0.2 µm s^−1^ quoted by Pollard and Borisy [46].

When the filament is stalled, *V* = 0, and the corresponding stall force *F*_0_ is given by:

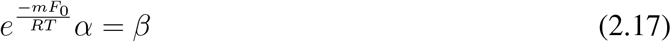

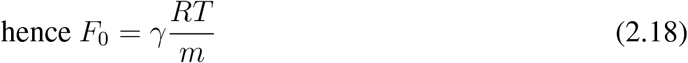

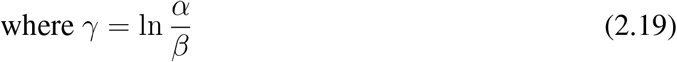

Using the values from Table 1:

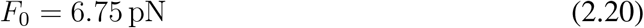

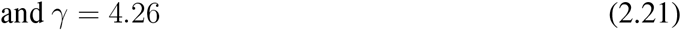

This value for *F*_0_ is within the range *F*_0_ = 7.7 *±* 1.3pN quoted by Abraham et al. [47, Table 2].

Moreover, using Equations (2.8),(2.9),(2.12),(2.13) and (2.19) the net reaction potential Φ_*C*_ is given as:

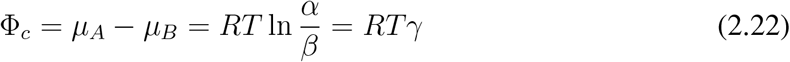

it follows that the stall force *F*_0_ can also be written as:

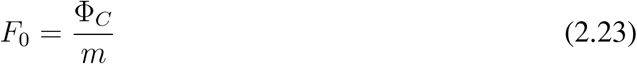

The force-velocity curve of Equation (2.14) can be simplified by using the expressions for *V*_0_ and *F*_0_ (2.15) and (2.18). In particular, equation (2.14) can be rewritten by normalising velocity *V* by *V*_0_ and force *F* by *F*_0_ to give:

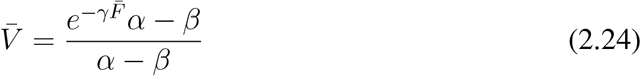

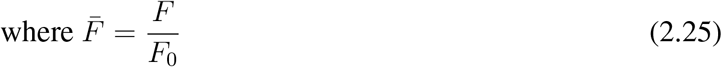

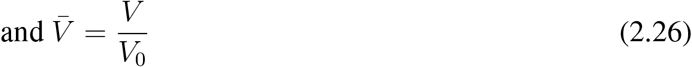

Equation (2.24) can be rewritten in terms of the single parameter *γ* (2.19) as:

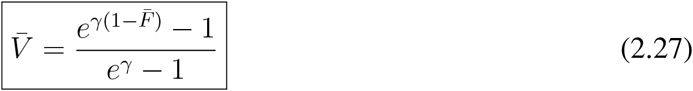

Figure 2(a) shows the variation of normalised mechanical velocity *V*_0_ with normalised mechanical force 0 ≤*F*_0_ ≤1 for three values of *γ*; the curve for *γ* = 4.26 corresponds to Figure 1 of Peskin et al. [32]. Figure 2(b) plots the same data as Figure 2(a) but in non-normalised form; this emphasises that both *F*_0_ and *V*_0_ increase with *γ*.

**Figure 2:**
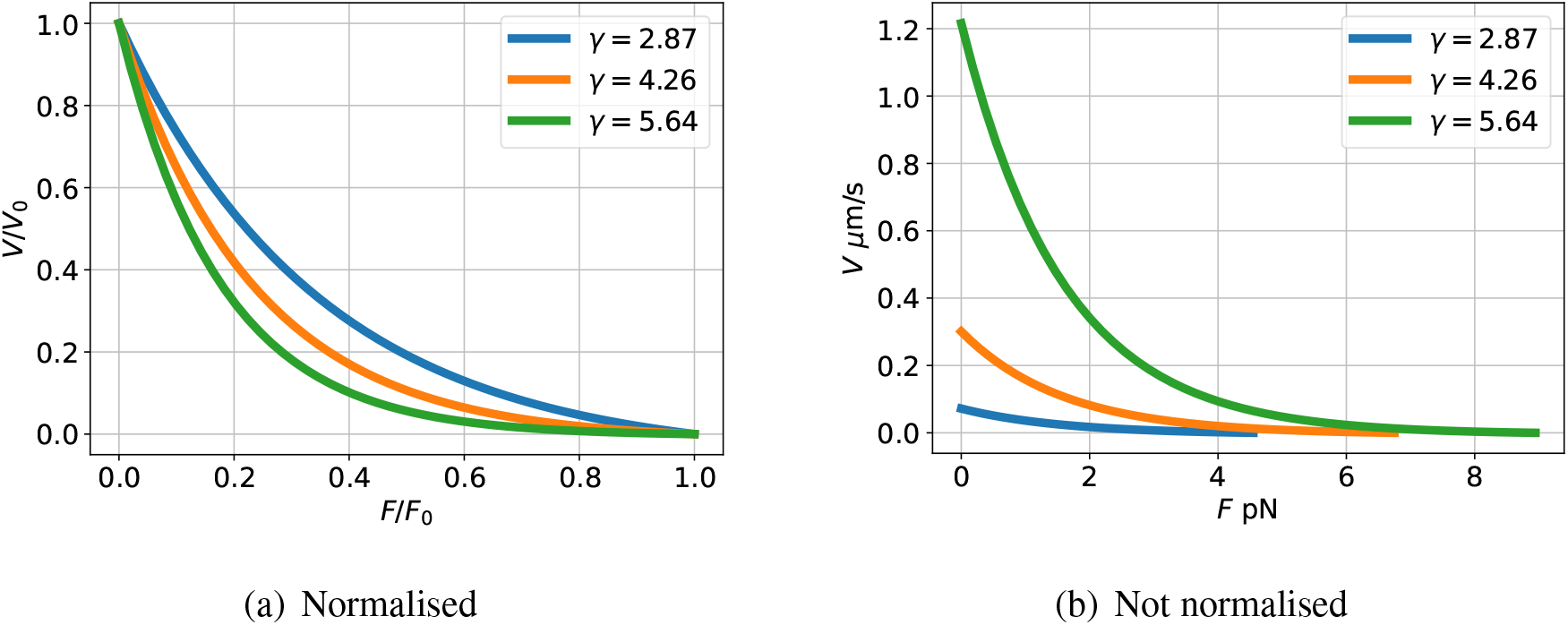
Force-velocity curve. Using the values from Table 1, *γ* = 4.26 and the transformer modulus *m* = 1.63 *×* 10^15^ m mol^−1^. The curves labelled *γ* = 4.26 are plotted for the values given in Table 1. The other two curves are for 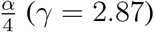 and 4*α* (*γ* = 5.64). (a) Normalised curves from Equation (2.27). (b) Without normalisation. This alternative representation emphasises that increasing the chemical potential *α* of the complex represented by **Ce**:**A** of Figure 1, and thus *γ* and the reaction potential Φ_0_, increases both the stall force *F*_0_ and the zero-force velocity *V*_0_.

## 3 Power and efficiency

The *efficiency* of living systems in general, and actin-based transduction in particular, is of great interest [34, 35, 48]. As the bond graph approach directly exhibits energy flow, it allows the power and the efficiency of the actin filament to be directly investigated.

The mechanical power *P*_*mech*_ (flowing though the **TF**) is:

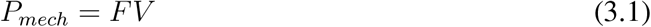

This is, of course equal to that part of the chemical power flowing into the **TF**. The chemical power *P*_*chem*_ into the system from the two complexes represented by **Ce**:**A** and **Ce**:**B** is the product of the molar flow *v* and the chemical reaction potential Φ_*c*_ (2.22):

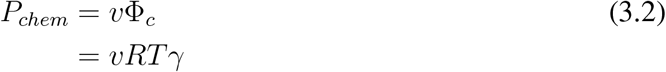

Using the values from § 2.3, *P*_*mech*_ and *P*_*chem*_ are plotted in Figure 3(a). There are many concepts of efficiency, one possibility is to define efficiency *η* as:

**Figure 3:**
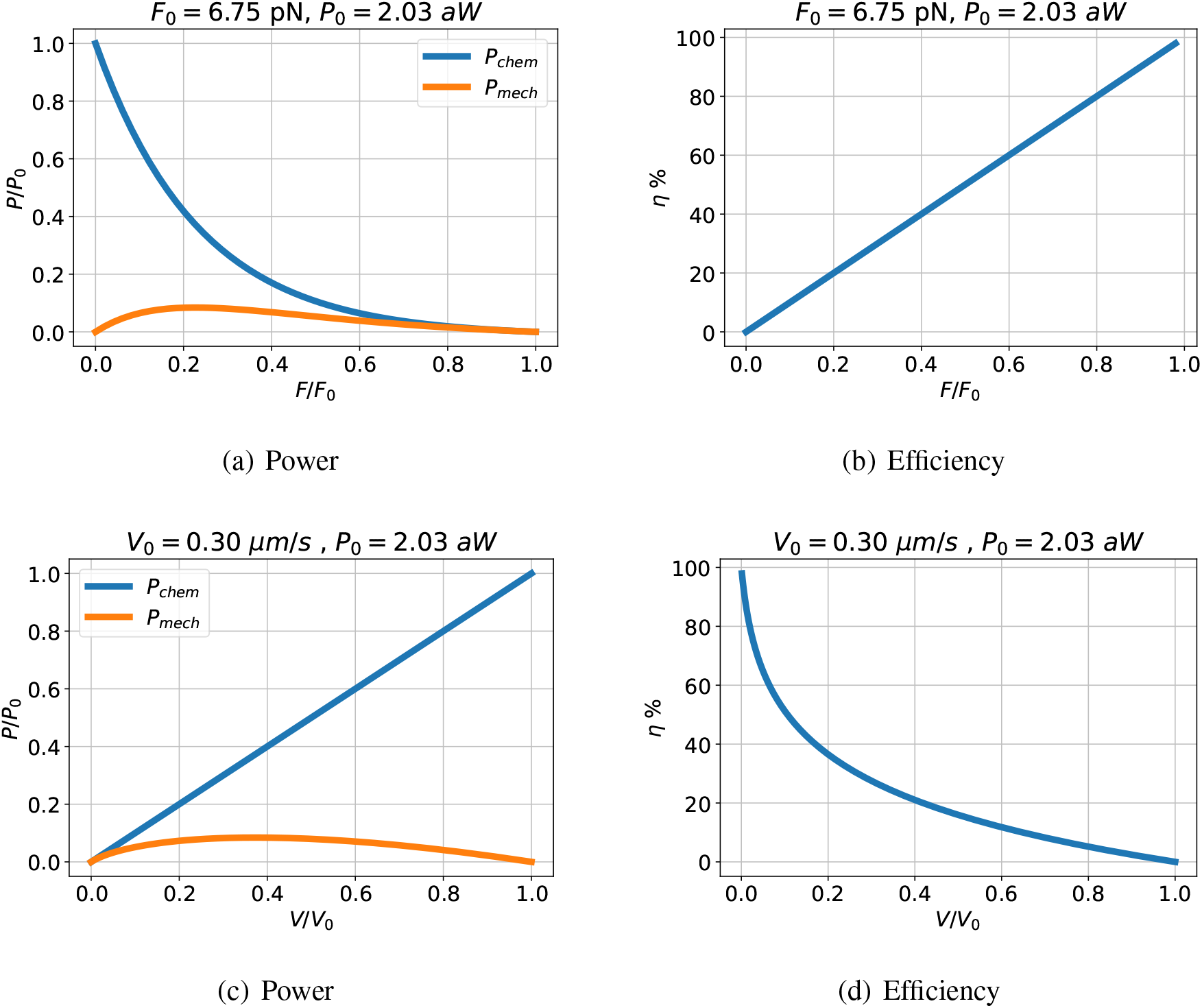
Power and Efficiency. (a) Using the values from Table 1 the normalised (by *P*_0_ = *F*_0_*V*_0_ = 2.03 aW) chemical power and mechanical power are plotted against normalised (by *F*_0_ = 6.75 pN) force. The chemical power reduces as force increases; the mechanical power (PV) is zero at two points: when the force *F* = 0 and the velocity *V* = 0 (corresponding to the stall force *F*_0_). The difference between the two powers is dissipated in the reaction component **Re**:**r**. (b) The *efficiency η* (3.3) is plotted against normalised force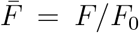; as indicated in Equation (3.4), the efficiency is equal to the normalised force. The efficiency is at its highest value when the system is in equilibrium: *V* = 0 and no energy is dissipated in the **Re** component. On the other hand, the highest power transfer occurs at *η* ≈ 20%; such is the tradeoff between power transfer and efficiency. (c) and (d). As (a) and (b) but plotted against normalised (by *V*_0_ = 0.30 µm) velocity.

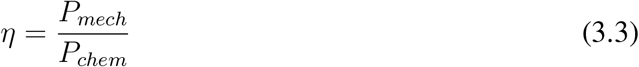

Using Equations (3.1), (3.2) and (2.23) is follows that efficiency *η* is proportional to normalised force:

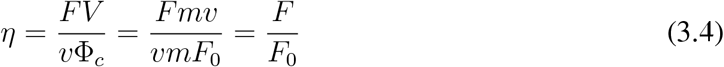

This is plotted in Figure 3(b). As expected, the efficiency is at its highest value when the system is in equilibrium: 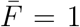 and 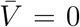 and no energy is dissipated in the **Re** component. On the other hand, the highest power transfer occurs at *η* ≈ 20%; such is the tradeoff between power transfer and efficiency.

## 4 Rigid filament not normal to surface

Mogilner and Oster [37] added two innovations to the model of Peskin et al. [32]: the actin rod impinges on the membrane at an angle *θ* to the normal (*θ* = 0 corresponds to normal to the membrane, 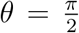 corresponds to parallel to the membrane); and the actin filament is flexible rather than rigid. The former is treated in this section, the latter in § 5. As discussed by Pollard [36] this work implies the existence of an optimal angle for maximum membrane velocity; this aspect is also examined in the bond graph context.

The rigid filament at an angle is shown in Figure 4(a) and the corresponding bond graph of Figure 4(b) is now derived. The bond graph approach models complex systems by incrementally adding more components. In the mechanical domain, the standard bond graph approach to modelling a coordinate rotation is to use an angle-modulated transformer **TF** component [7–9, 49]. With reference to Figure 4, the transformer **TF**:**c** has a modulus cos(*θ*) and transforms *F* and *V* in rod coordinates to *F*_*m*_ and *V*_*m*_ in membrane coordinates. In particular:

**Figure 4:**
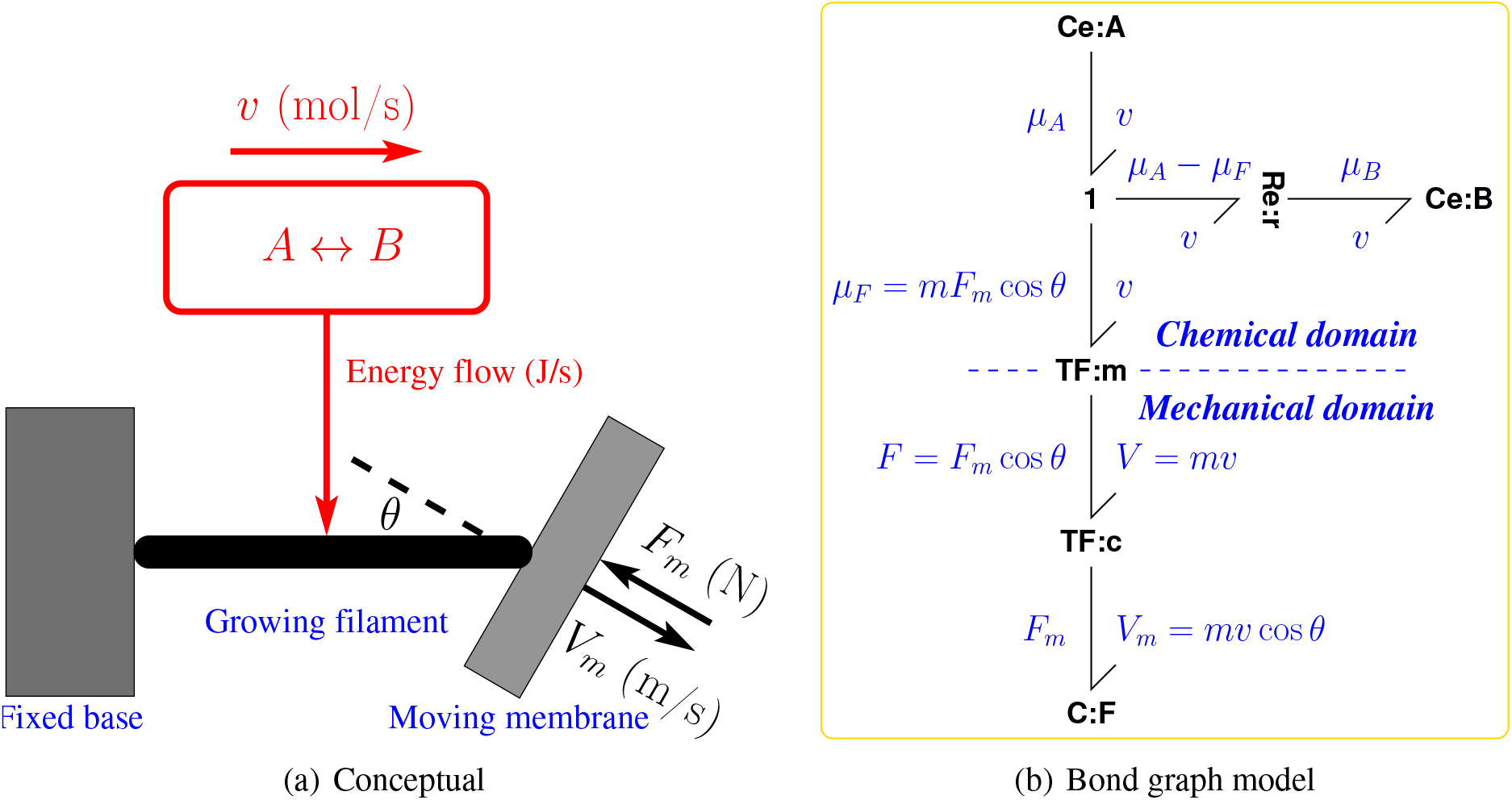
Bond Graph model – rigid filament at angle *θ* ≠ 0 with normal to surface. (a) This is identical to Figure 1(a) except that the growing filament is at an angle of *θ*rad to the membrane normal. As discussed in the text, the filament force *F* and velocity *V* are now different to the membrane force *F*_*m*_ and velocity *V*_*m*_. (b) This is identical to Figure 1(b) except that the additional **TF** component **TF**:**c** is used to perform the geometric transformation between *F, V* and *F*_*m*_,*V*_*m*_ imposed by the angle *θ* of Equations (4.1)& (4.2). It is assumed that the actin filament is in contact with a smooth surface and therefore produces no force parallel to the surface.

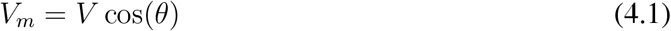

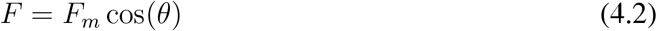

Substituting Equations (4.1) and (4.2) into Equation (2.27) gives the formula giving normalised membrane velocity 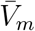 in terms of normalised membrane force 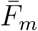 with the parameters *γ* and *θ*:

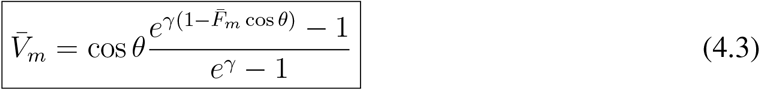

where

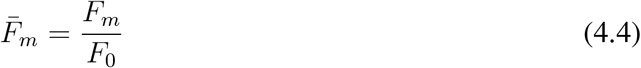

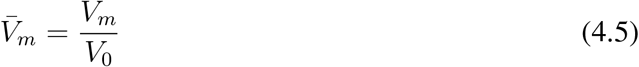

Using the same parameters as for Figure 2, and plotting normalised membrane velocity *V*_*m*_ against *θ* for a range of values of force *F*_*m*_ gives the curves of Figure 5(a); the vertical lines locate the optimal angle (giving the maximum membrane velocity) for each value of force. The optimum value of *θ* plotted against force is given in Figure 5(c). Note that it is possible to get forward membrane velocity for force greater than the zero stall force *F*_0_ for *θ >* 0. Indeed, the effective stall force is *F*_0_*/* cos *θ*; this is plotted in Figure 5(d).

**Figure 5:**
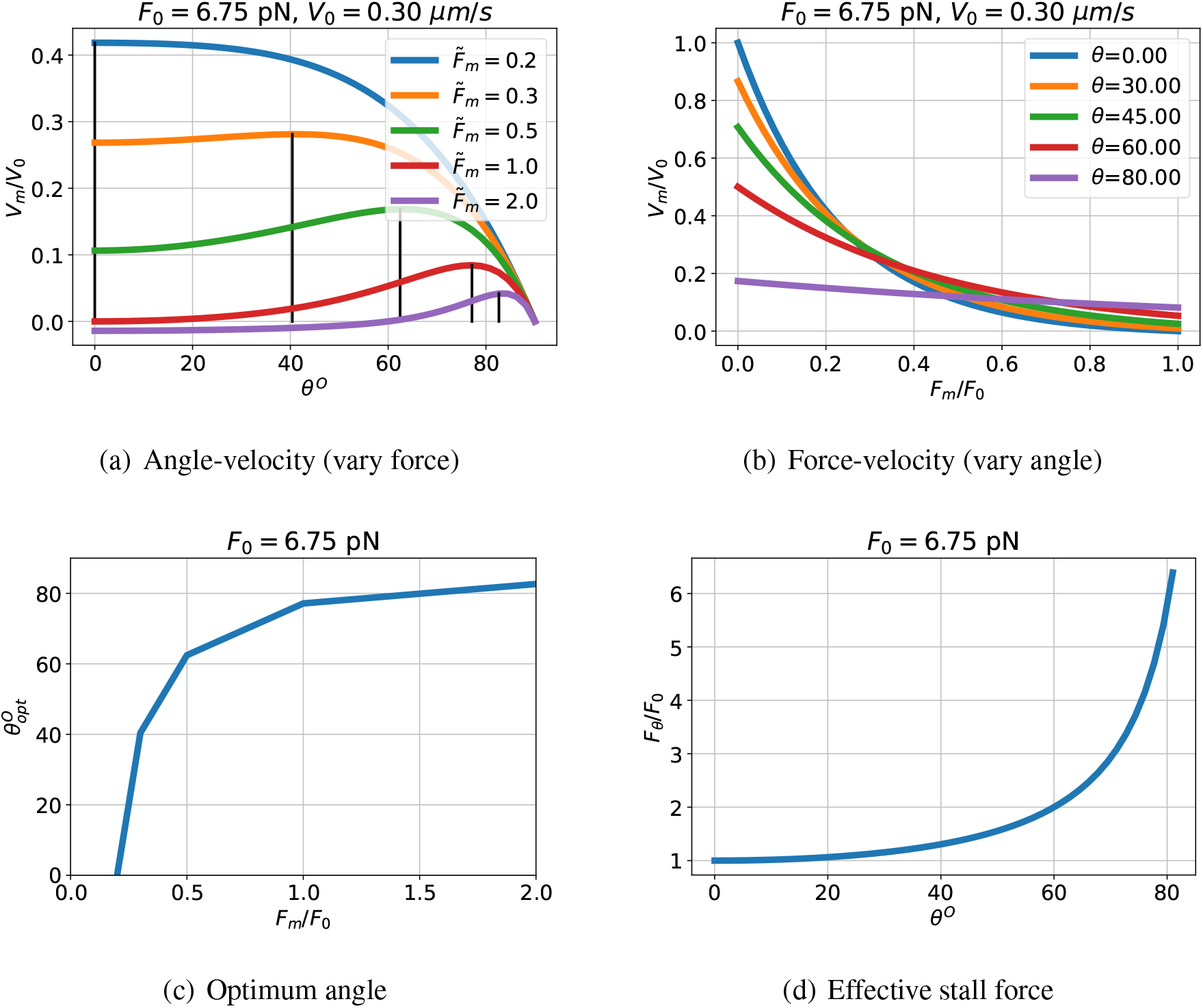
Non-normal angle: *θ* ≠ 0: properties. (a) The normalised membrane velocity 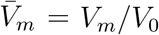 is plotted against incidence angle *θ* for various values of the normalised membrane force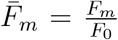. The optimal incidence angle for each value of force is marked. (b) The normalised membrane velocity 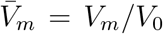 is plotted against normalised membrane force 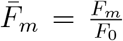 for various values of the incidence angle *θ*. (c) The optimal incidence angle *θ*_*opt*_ is a function of membrane force *F*_*m*_. (d) The effective stall force *F*_*θ*_ is a function of incidence angle *θ*.

## 5 Flexible filament not normal to surface

If the actin filament is flexible, it will bend if a torque *τ* is applied; this could be analysed using standard beam theory [50]; but, as a first approximation, the flexible beam is replaced by a rigid beam of length *L* connected to the base by a rotational spring with compliance *σ* radians*/*Nm and angle of rotation *ϵ*. Figure 6(a) shows where the tip impinges on the membrane. The angle between the membrane and fixed base is *θ*_0_ and so the angle *θ* between filament and membrane normal is given by:

**Figure 6:**
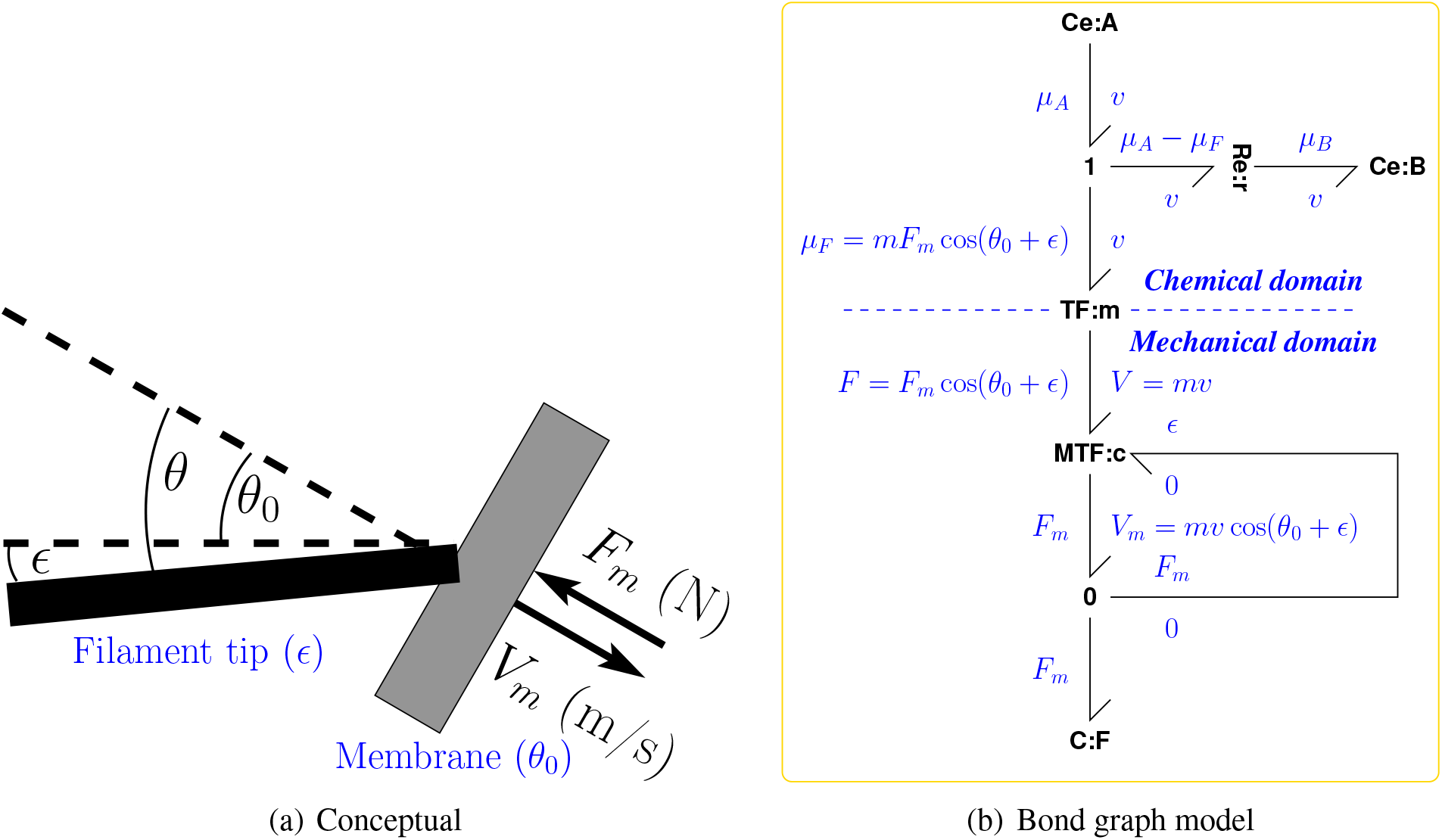
Bond Graph model – flexible filament. (a) This is identical to Figure 4(a) except that only the growing tip is shown, the growing filament is assumed to be flexible and the flexibility approximated by a spring with compliance *σ* (N m rad^−1^) at an angle of *ϵ* (rad) to the base normal. The membrane is tilted by an angle *θ*_0_ and the angle between filament and membrane normal is *θ* = *θ*_0_ + *ϵ*. (b) This is identical to Figure 4(b) except that the **TF** component **TF**:**c** is replaced by a *modulated* transformer **MTF**:**c** modulated by the flexibility angle *ϵ* as generated by Equation (5.10) from the membrane force *F*_*m*_. The **0** junction ensures that the effort (force *F*) on the three impinging bonds is identical.

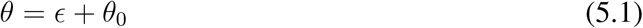

Substituting for *θ* in Equation (4.3) gives:

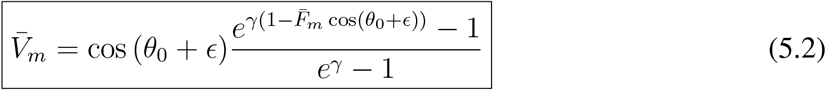

*ϵ* can be computed as follows. Using the equations for a linear simple spring with compliance *σ*:

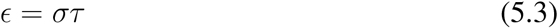

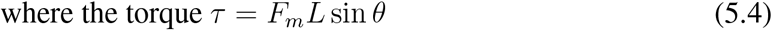

Combining Equations (5.1) – (5.4):

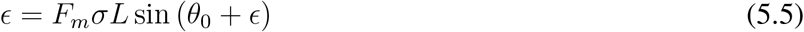

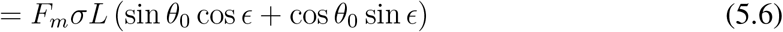

Hence

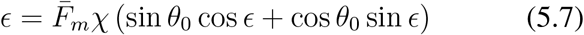

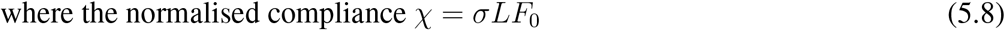

and 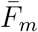 is given by (4.4). Following Mogilner and Oster [37, Appendix B], it assumed that *ϵ* ≪ *π*/2 and thus Equation (5.7) becomes:

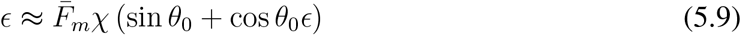

hence:

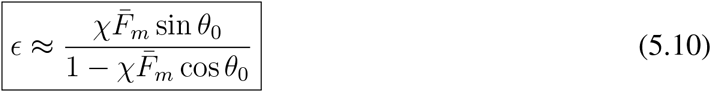

The corresponding bond graph is given in Figure 6(b). The modulated transformer **MTF** [49] allows the **TF** component to be modulated by *ϵ* (5.1) generated from the membrane force *F*_*m*_ using Equation (5.10).

The velocity-force curve is given by Equations (4.3), (5.1) and (5.10). Figure 7 summarises solutions to these equations for χ = 0.47 (see § 5.1, Equation (5.17)) which can be compared to the rigid case of § 4 and Figure 5(d).

**Figure 7:**
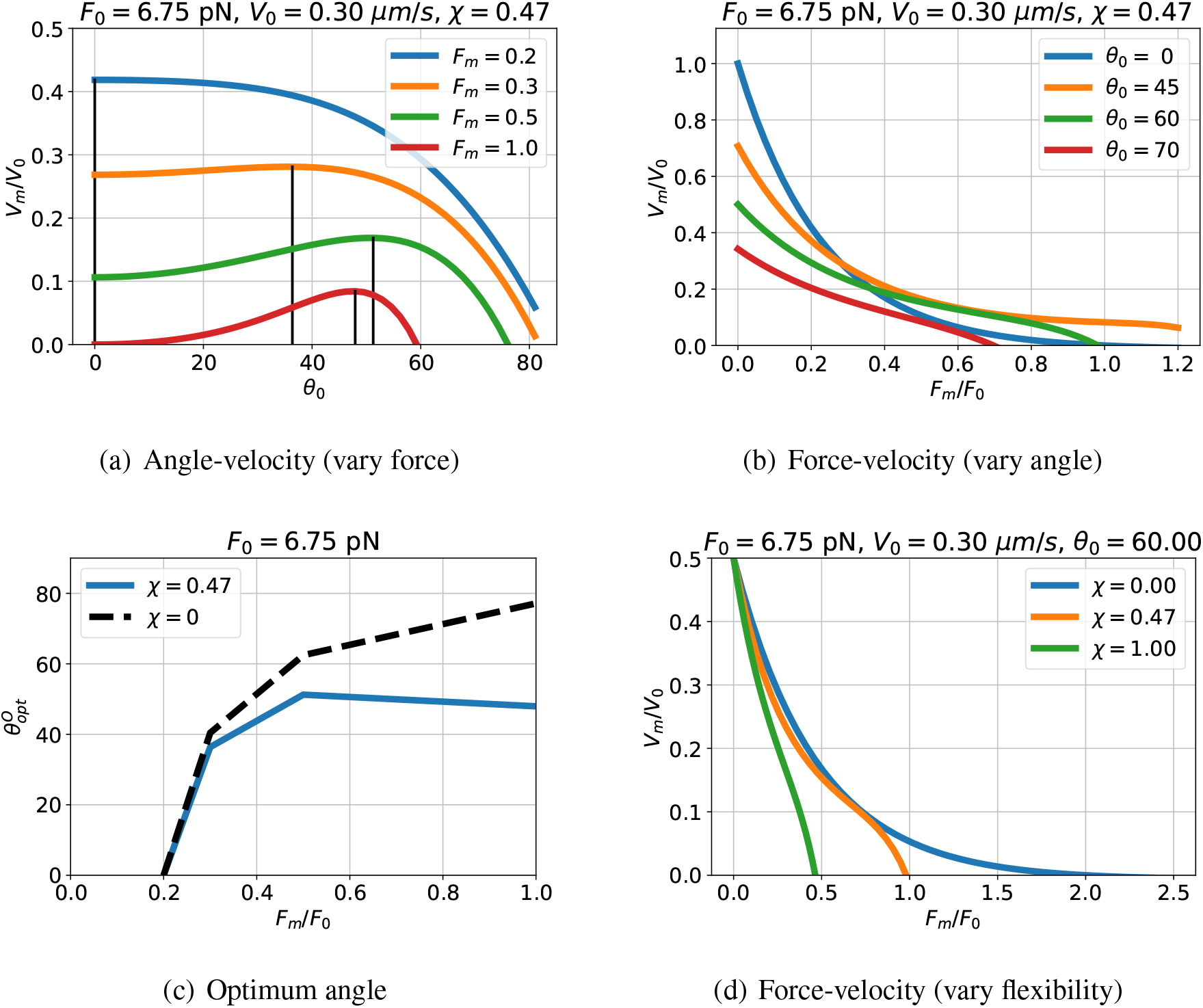
Flexible filament. (a) Force/velocity diagram for various values of angle *θ*_0_ and χ = 0.47 (see § 5.1, Equation (5.17)). (b) Force/velocity diagram for various values of normalised compliance χ and *θ*_0_ = 60°. (c) The optimum (maximum velocity) values of *θ* are plotted for the flexible (χ = 0.47) and rigid (χ = 0) cases. (d) The velocity force diagram for the flexible filament; this should be compared to the rigid filament result of Figure 5(a).

### 5.1 Computation of filament normalised compliance χ

The formula (5.10) contains the normalised compliance χ a a parameter. This section computes the value of χ corresponding to the parameters listed in Table 1.

Mogilner and Oster [37] use the *persistence length λ* as a measure of filament stiffness where:

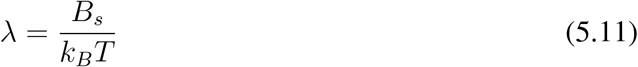

where *B*_*s*_ is the filament *bending stiffness* (*B*_*s*_ = *EI* in beam theory), *k*_*B*_ is the Boltzmann constant and *T* the absolute temperature. Modelling the filament as an Euler cantilever beam of length *L* with a force *F*_*T*_ at the tip, the deflection *δ* is [50, § 21, Chapter 3]:

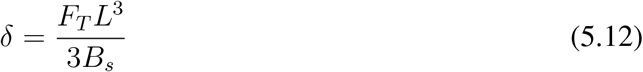

Thus the deflection angle *ϵ* with respect to the root is:

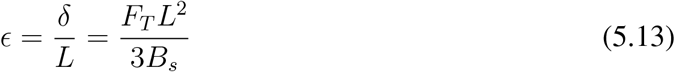

Noting that the torque *τ* = *F*_*T*_ *L*, the equivalent compliance *σ* is given by:

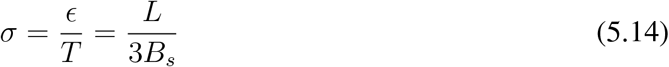

Using equation (5.11), the equivalent compliance *σ* becomes:

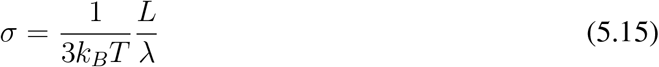

hence, using equation (5.8):

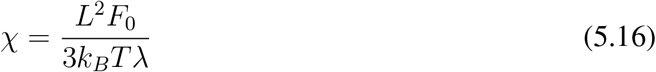

Using the parameters from Table 1:

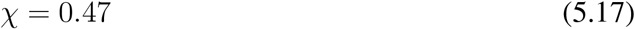

## 6 Curvature of the force-velocity curve

The force-velocity curves of Figure 2 are *convex* in the sense that the second derivative is positive. A number of authors [41–44] have shown that the effect of multiple filaments interacting with the properties of the membrane can lead to force velocity curves which are entirely or partially concave. This section shows that the effect of the filament compliance discussed in § 5 can lead to a similar effect.

Figure 7(d) shows force-velocity curves for varying compliance. Although when normalised compliance χ = 0 the curve is convex, the curves for *χ >* 0 become concave as force increases. Figure 8 examines this behaviour further.

**Figure 8:**
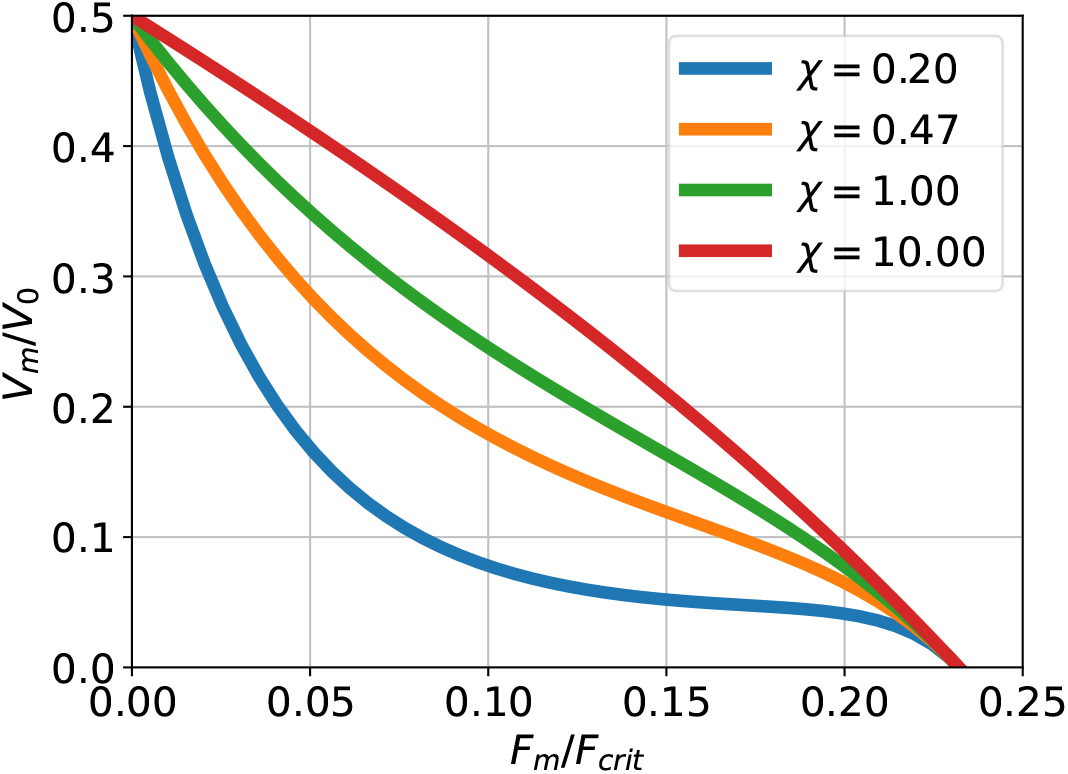
Curvature of the force-velocity curve. Figure 7(d) is replotted with a new range of values of normalised compliance χ. Moreover, the force is normalised by the critical force *F*_*crit*_ (6.1) instead of the stall force *F*_0_ (2.18) to emphasise how the curvature varies with the normalised compliance χ. Although (from Figure 7(d)) the force-velocity curve is convex when the compliance is zero χ = 0 part of the curve becomes concave as χ increases.

Equation (5.10) exhibits a singularity at 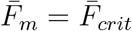 where

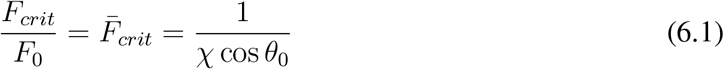

Figure 8 is identical to Figure 7(d) except that the force is normalised by *F*_*crit*_ rather than *F*_0_ and it is plotted for a range of *χ >* 0. Thus for small normalised compliance (χ = 0.2) the curve is convex for small values of normalised force *F*_*m*_*/F*_*crit*_; the curve is entirely concave for large normalised compliance (χ = 10). As discussed previously in the multi-filament context [51] such a concave curve leads to an initial plateau in the force-velocity curve giving lower sensitivity of velocity to force in the region.

## 7 Experimental comparison

Li et al. [35, Fig. 1D] provide experimental values of actin growth velocity for various values of growth stress obtained from branched actin network impinging on a cantilever. As they point out, force-feedback occurs in that the density of actin filaments increases with stress and this also appears in [35, Fig. 1D]. For the purposes of this paper, the behaviour of an average filament is obtained by normalising experimental pressure by the measured actin density^3^.

This section fits the velocity-force curve for the flexible filament of § 5 given by Equations (5.2) & (5.10) to the normalised experimental data. There are 5 parameters: the angle *θ*_0_, the chemical driving force encapsulated in *γ* (2.19), the normalised compliance χ (5.8), the stall force *F*_0_ (2.18) and the unloaded velocity *V*_0_ (2.15).

Li et al. [35] give the filament contact angle as 54°. As the contact angle is relative to the surface (as opposed to the surface normal); this corresponds to *θ*_0_ = 90 − 54 = 36°. *F*_0_ and *V*_0_ were approximated by the maximum values of the normalised force and velocity respectively.

The remaining two parameters *γ* and χ were estimated by fitting the velocity-force curve for the flexible filament of § 5 given by Equations (5.2) & (5.10) to the normalised experimental data using the Python minimisation routine `scipy.optimize.minimize` using the default BFGS (Broyden–Fletcher–Goldfarb–Shanno) [52] method. This gives *γ* = 3.15 and χ = 0.29; in comparison, the values of *γ* = 4.26 from Equation (2.21) and χ = 0.47 from Equation (5.17) correspond to Table 1 and were used in Figure 7. The corresponding curve is plotted as the firm line in Figure 9 with the experimental points superimposed.

**Figure 9:**
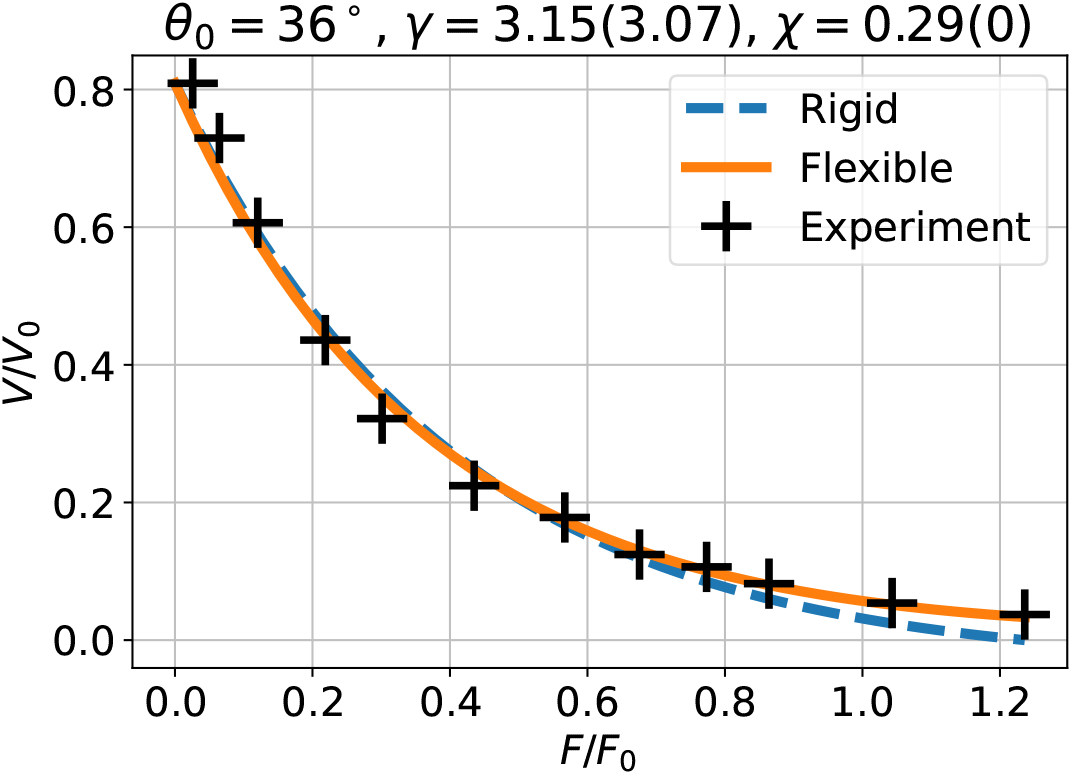
Experimental comparison of the formulae (5.2) & (5.10) with experimental data presented by Li et al. [35] based on results of Bieling et al. [34]. The data is extracted from the additional information in elife-73145-fig1-data1-v2.xlsx. The value of *θ*_0_ = 36° is taken from Li et al. [35]. Using the rigid model (§ 4, dashed line), the parameter *γ* (2.19) (the chemical driving force) is estimated. The estimated value is *γ* = 3.07. Using the flexible model (§ 5, firm line), the two parameters are estimated: *γ* and the normalised compliance χ (5.8). The estimated values are *γ* = 3.15 and χ = 0.29.

For comparison, the data was fitted with the rigid model (χ = 0) by estimating the single parameter *γ*. This is shown as the dashed line in Figure 9. The deviation from experimental data is greater as normalised force increases; this corresponds to the greater effect of flexibility at larger forces. The deviation would increase still further for larger values of membrane force *F*_*m*_.

## 8 Conclusion

A novel bond graph approach to the modelling of chemomechanical transduction has been presented using single-filament actin polymerisation as an example. Despite its relative simplicity, the bond graph approach has been shown to give an identical formula to the Brownian ratchet approach presented in the seminal paper of Peskin et al. [32]. The fact that two apparently disparate approaches give the same result seems to be due to the fact that bond graphs for chemical systems are, like the Brownian ratchet, built on thermodynamic principles [10, 11]. It is of historical interest to note that Oster was involved in establishing the foundations of not only the energy-based chemical bond graph [10, 11] but also the Brownian ratchet [32, 37].

The approach in this paper is closely related to previous work [19, 21] on the bond graph representation of chemoelectrical transduction where the **TF** component is used to link chemical and electrical domains using a modulus *m* = *z***F** where *z* is the number of electrical charges and **F** (C mol^−1^) the Faraday constant. Indeed, the stall force formula (2.18) relating the chemical and mechanical domains is analogous to Nernst’s equation relating the chemical and electrical domains.

Although, as emphasised by Peskin et al. [32] and Pollard [36], energy-based approaches “provide no mechanistic explanation of how the free energy of polymerisation is actually transduced into directed mechanical force”, the bond graph approach has the advantage that a model of an actin filament can be readily extended by adding additional components or modules. For example, it was shown in his paper that the basic model of actin chemomechanical transduction can be extended to non-normal incidence and flexible filaments by simply adding more components to the bond graph representation. More generally, the bond graph approach is inherently modular [13, 16, 17], and bond graph modules containing both basic bond graph components and modules can be constructed in a hierarchical fashion. In this context, future work will investigate encapsulating the bond graph of actin chemomechanical transduction as a bond graph module and using the modular bond graph approach to incrementally build models of more complex systems including networks of actin filaments, adhesion, capping, moving elastic membranes and actin recycling. The actin chemomechanical transduction is driven by ATP hydrolysis; this could be explicitly linked to bond graph models of metabolism. Using the approach of § 3, efficiency at the cellular level could then be investigated.

Understanding the flow of cellular energy is crucial to understanding life [48]; therefore energy-based bond graph modelling including the chemical, electrical and mechanical domains provides a route to this understanding.

## Supporting information

A Short Introduction to Bond Graph Modelling

## 9 Author Contributions

PJG prepared the numerical results and drafted the manuscript; PJG, MP and VR jointly developed the concepts, discussed the results and revised the manuscript. All authors approved the final version for publication.

## 10 Acknowledgements

PJG would like to thank the Faculty of Engineering and Information Technology, University of Melbourne, for its support via a Professorial Fellowship. The authors would like to thank the anonymous reviewers for their constructive comments which have enhanced the paper.

## Footnotes

1 The notion of a chemical complex as the combination of a number of chemical species arises from Chemical Reaction Network Theory [45]) and is discussed in the Supplementary Material.

2 See the Supplementary material for more details

3 Interpolation and extrapolation using a quadratic curve fit was used to create values of actin density for the values of experimental pressure in the velocity dataset.

