## Supplementary material for "Energy-based Modelling of Single Actin Filament Polymerisation Using Bond Graphs": A Short Introduction to Bond Graph Modelling

September 2, 2024

**Abstract**

The purpose of this document is to provide the bond graph background necessary to understand the paper itself. More detail about general bond graph theory is to be found in references [1–3] and more detail about biochemical systems in references [4–10].

### Contents

**S1 A Short Introduction to Bond Graph Modelling** **S-1**
S1.1 Basic concepts . . . . . S-1
S1.2 Example: an electrical system . . . . . S-2
S1.3 Example: a chemical system . . . . . S-3
S1.3.1 Complexes . . . . . S-4
S1.4 Example: a mechanical system . . . . . S-4
S1.5 Connecting systems: the **TF** component . . . . . S-5

### S1 A Short Introduction to Bond Graph Modelling

The purpose of this section is to provide the bond graph background necessary to understand the paper itself. More detail about general bond graph theory is to be found in references [1–3] and more detail about biochemical systems in references [4–10].

#### S1.1 Basic concepts

| Bond Graph | Electrical | Mechanical | Chemical |
|---|---|---|---|
| Effort<br>$e$ | Voltage<br>$\phi$ (V) | Force<br>$F$ (N) | Gibbs energy<br>$\mu$ ($\mathrm{J\,mol^{-1}}$) |
| Flow<br>$f$ | Current<br>$I$ (A) | Velocity<br>$V$ ($\mathrm{m\,s^{-1}}$) | Molar flow<br>$v$ ($\mathrm{mol\,s^{-1}}$) |
| Quantity<br>$q = \int^t f(\tau)d\tau$ | Charge<br>$Q$ (C) | Displacement<br>$X$ (m) | Molar amount<br>$x$ (mol) |

Table S1: Analogous variables. Systematic modelling, including the bond graph approach, uses the concept of analogous variables to bring together different physical domains. One such analogy is the effort/flow analogy displayed here: each row contains analogous variables, each column corresponds to a domain. In each case effort $\times$ flow $=$ power. Gibbs energy is also referred to as *chemical potential*. Other physical domains such as magnetic and hydraulic can be represented in a similar fashion.

Bond graphs unify the modelling of energy within and across multiple physical domains using the concepts of *effort* ($e$) and *flow* ($f$) variables whose product is *power*. As indicated in Table S1, in the electrical domain the effort variable is voltage (electrical potential) $\phi$ with units of $\mathrm{V}$ and the flow variable is current $i$ with units of $\mathrm{A}$ (or $\mathrm{C\,s^{-1}}$) which have the product $P = \phi i$ with units of $\mathrm{J\,s^{-1}}$ or $\mathrm{W}$. In the mechanical domain, the effort variable is Force $F$ with units of $\mathrm{N}$ and the flow variable is velocity $V$ with units of $\mathrm{m\,s^{-1}}$ have the product $P = FV$ with units of $\mathrm{J\,s^{-1}}$. In the chemical domain, the effort variable is *chemical potential* $\mu$ with units of $\mathrm{J/mol}$ and the flow variable is reaction flow rate $v$ with units of $\mathrm{mol/sec}$; $P = \mu v$ also has units of $\mathrm{J\,s^{-1}}$.

The bond graph **C** component is a generic potential energy *storage* component corresponding to a *capacitor* in the electrical domain, a *spring* in the mechanical domain and the accumulation of a chemical species in the chemical domain. The electrical capacitor generates voltage (electrical potential) $\phi$ according to the *linear* relationship: $\phi = Q/C$ where $C$ is the capacitance in Farads (F) and $Q$ the charge in Coulombs (C) . Similarly, a linear mechanical spring generates force $F$ according to the *linear* relationship: $F = K_s X$ where $K_s$ is the spring constant ($\mathrm{N\,m^{-1}}$) and $X$ the displacement in metres.

In contrast, the chemical species generates chemical potential $\mu \mathrm{J\,mol^{-1}}$ according to the *nonlinear* relationship:

$$\mu = RT \ln Kx \qquad \text{(S1.1)}$$

where $R$ is the universal gas constant with units of $\mathrm{J\,mol^{-1}\,K^{-1}}$, $T$ absolute temperature with units of $\mathrm{K}$, $x = q/q_0$ where $q_0$ is the reference quantity and and $K$ a species-dependant constant which is dimensionless and depends on the reference quantity. Because of the special nonlinear relationship, the chemical version of the **C** component is given a special symbol: **Ce** .

The bond graph **R** component is a generic energy *dissipation* component corresponding to a *resistor* in the electrical domain and a damper in the mechanical domain. In the electrical domain, it generates current $i$ according to the *linear* relationship: $i = \kappa\Delta\phi$ where $\kappa$ is the conductance and $\Delta\phi$ is the net potential across the resistor. Similarly, a linear mechanical damper generates force $F$ according to the *linear* relationship: $F = d\Delta V$ where $d$ is the damper constant $(\mathrm{N\,m^{-1}\,s})$ and $\Delta V$ the relative velocity $(\mathrm{m\,s^{-1}})$ between the damper terminals.

The **R** component corresponds a chemical *reaction* in the chemical domain. Assuming mass-action kinetics it generates molar flow $v$ according to the *nonlinear* relationship:

$$v = \kappa\left(\exp\frac{A^f}{RT} - \exp\frac{A^r}{RT}\right) \tag{S1.2}$$

where $A^f$ (the *forward affinity*) and $A^r$ (the *reverse affinity*) are the net chemical potentials (with units of $\mathrm{J\,mol^{-1}}$) of the reactants and products of the reaction respectively; $\kappa$ is the reaction *rate constant* with units of $\mathrm{mol\,s^{-1}}$. Because this relationship treats the forward and reverse affinities differently, the reaction version of the **R** component is given a special symbol: **Re** .

It can be convenient to rewrite Equation (S1.2) as [4]:

$$v = \kappa\left(v_0^+ - v_0^-\right) \tag{S1.3}$$

$$\text{where } v_0^+ = \exp\frac{A^f}{RT} \tag{S1.4}$$

$$v_0^- = \exp\frac{A^r}{RT} \tag{S1.5}$$

The bond graph **I** component is a generic kinetic energy *storage* component corresponding to a *inductance* in the electrical domain, a *mass* in the mechanical domain - there is no analogue in the chemical domain. The **I** component is not used in the paper but appears in § S1.4.

Bond graphs represent the flow of energy between components by the bond symbol $\rightharpoonup$ where the direction of the harpoon corresponds to the direction of positive energy flow; this is a sign convention. The bonds connect components via *junctions* which transmit, but do not store or dissipate energy. There are two junctions: the **0** junction where all impinging bonds have the same *effort* and the **1** junction where all impinging bonds have the same *flow*. The expression for the flows associated with the **0** junction, and the efforts associated with the **1** junction, are determined by the energy conservation requirement. A further energy transmission component is the *transformer* (**TF** ) component discussed in § S1.5.

#### S1.2 Example: an electrical system

Figure S1 illustrates bond graph modelling in the electrical domain. The four capacitors (A–D) and two resistors ($r_1$,$r_2$) are connected as in the schematic diagram of Figure 1(a). In bond

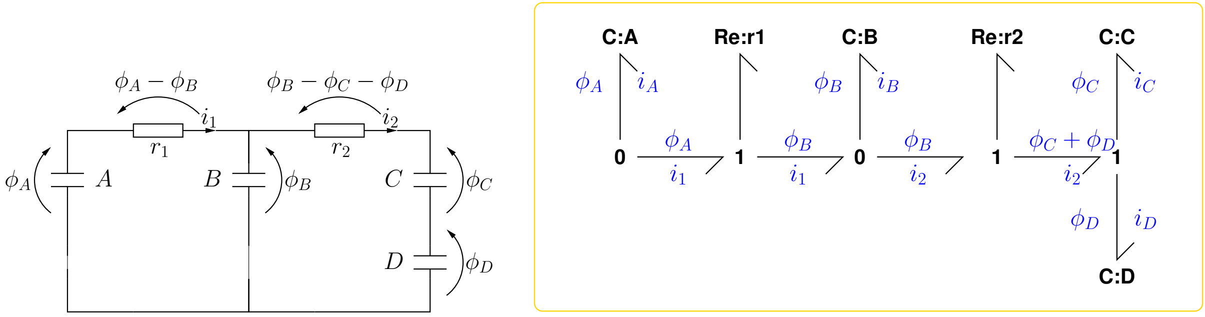


(a) Schematic (b) Bond Graph

Figure S1: Modelling Electrical Systems. (a) Electrical schematic diagram. (b) Bond graph: $\Delta\phi_1 = \phi_A - \phi_B$, $\Delta\phi_2 = \phi_B - (\phi_C + \phi_D)$.

graph colon notation, the symbol before the colon indicates the component type and the symbol after the colon indicates the component name. Thus the bond graph of Figure 1(b) has four **C** components named A–D to represent the four capacitors and a two **R** components named $r_1$ & $r_2$ to represent the resistors. The **0** junctions impose a common potential on the attached bonds (parallel connection) The **1** junctions impose a common current (flow) on the attached bonds (series connection) Given the constitutive relations for the **C** and **R** component, the dynamical equations describing the system can be automatically generated from the bond graph of Figure 1(b).

#### S1.3 Example: a chemical system

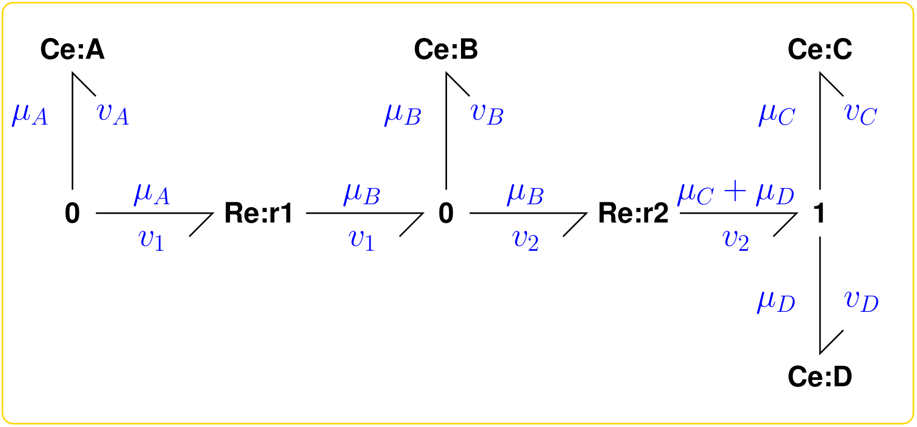


Figure S2: Modelling Chemical Systems. $A \overset{r1}{\rightleftharpoons} B \overset{r2}{\rightleftharpoons} C + D$. $v_A = -v_1$, $v_B = v_1$,$v_C = v_D = v_2$.

Figure S2 illustrates bond graph modelling in the chemical domain; the corresponding reaction is $A \overset{r1}{\rightleftharpoons} B \overset{r2}{\rightleftharpoons} C + D$. In a similar manner to the electrical system of Figure S1, the bond graph of Figure S2 has four **C** components named A–D to represent the four species and a two **Re** component named r1 and r2 to represent the two reactions. The bond graphs of the electrical and chemical system are similar; the major differences are the use of the **Ce** component in place of the **C** component and **Re** component to replace the **1** & **R** combinations of Figure 1(b). These differences are discussed in § S1.1.

The affinities (net potentials) are:

$$A_1^f = \mu_A \tag{S1.6}$$

$$A_1^r = \mu_B \tag{S1.7}$$

$$A_2^f = \mu_B \tag{S1.8}$$

$$A_2^r = \mu_C + \mu_D \tag{S1.9}$$

Using Equations (S1.1) & (S1.2):

$$v_1 = \kappa_1 \left( \exp \frac{A_1^f}{RT} - \exp \frac{A_1^r}{RT} \right) = \kappa_1 \left( K_A x_A - K_B x_B \right) \tag{S1.10}$$

$$v_2 = \kappa_2 \left( \exp \frac{A_2^f}{RT} - \exp \frac{A_2^r}{RT} \right) = \kappa_2 \left( K_B x_B - K_C K_D x_C x_D \right) \tag{S1.11}$$

##### S1.3.1 Complexes

Chemical Reaction Network Theory provides a means of representing a network of chemical reactions in terms of a (mathematical) graph [11]. A key to this is the notion of *complexes*. In the context of this simple introduction, complexes are exemplified by the reaction network of Figure S2: there are two reactions $r_1$ and $r_2$ and three complexes: A, B & C + D. In other words, the complexes are the species, or sum of species, appearing on one side or other of a reaction. In this case, the formulation of Equation (S1.3) is particularly useful as the definition of $v_0^+$ and $v_0^-$ is not dependent on the the number of species in each complex but rather just the net affinities $A^f$ and $A^r$.

Further details of the relationship between complexes and bond graphs are given by Gawthrop and Crampin [4] and by Gawthrop and Pan [12].

#### S1.4 Example: a mechanical system

Figure S3 illustrates bond graph modelling in the mechanical domain using the standard mass-spring-damper system of Figure 3(a). The bond graph of Figure 3(b) represents the mass, spring and damper by the **I** , **C** and **R** components described in § S1.1.

The **1** junctions impose a common velocity (flow) on the attached bonds; the **0** junction is not strictly necessary but could be used to attach further components or systems. Again, given the constitutive relations for the **I** , **C** and **R** components, the dynamical equations describing the system can be automatically generated from the bond graph of Figure 3(b).

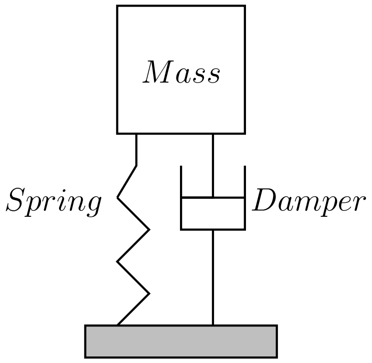


(a) Schematic

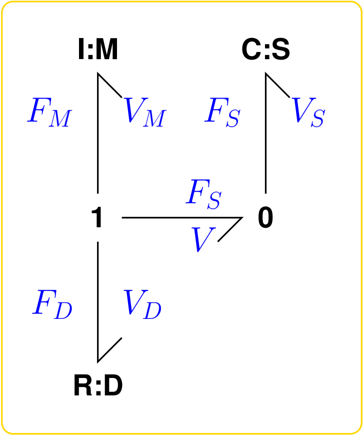


(b) Bond Graph

Figure S3: Modelling Mechanical Systems. (a) Schematic diagram. The system comprises a mass, a spring and a damper attached to a rigid base. The three components share a common velocity $V_S = V_M = V_D = V$. (b) Bond graph. The junction structure ensures that the three velocities are equal, and that the three forces sum to zero: $F_S + F_M + F_D = 0$.

#### S1.5 Connecting systems: the **TF** component

The **TF** component transmits power without dissipation or storage. It has two main uses: modelling power transmission *within* a physical domain and modelling power transduction *between* domains. In the first case, for example, a mechanical lever can be modelled as the **TF** of Figure 4(a) and chemical stoichiometry can be modelled as the **TF** of Figure 4(b). In the second case, chemoelectrical transduction can be modelled as the **TF** of Figure 4(c) making use of Faraday's constant which converts molar flow $(\mathrm{mol\,s^{-1}})$ to electron flow $(\mathrm{C\,s^{-1}})$.

Figure 4(d) lies at the heart of the paper: molar flow $(\mathrm{mol\,s^{-1}})$ is converted to the filament tip velocity $(\mathrm{m\,s^{-1}})$.

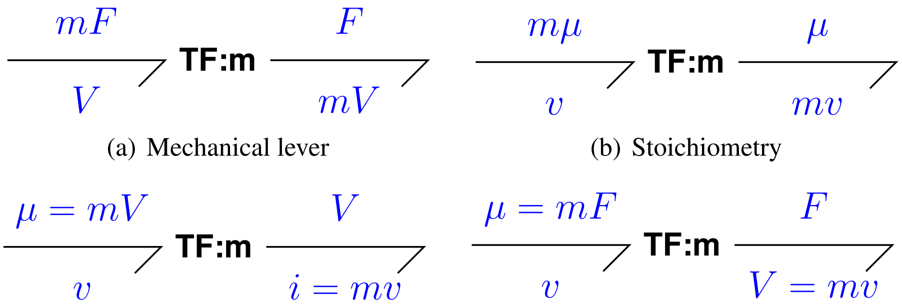


(a) Mechanical lever (b) Stoichiometry (c) Chemoelectrical transduction (d) Chemomechanical transduction

Figure S4: The **TF** component. In each case the power (product of effort and flow) at the two ports is the same. (a) Mechanical lever: the modulus $m$ is the ratio of the velocities at each end of the lever [1, 2]. (b) Chemical stoichiometry: the modulus $m$ is the ratio of the flows at the two ports; it is usually integer [12]. (c) Chemoelectrical transduction:the modulus $m$ is Faraday's constant which has units $(\mathrm{C\,mol^{-1}})$ [13, 14]. (d) Chemomechanical transduction: the modulus $m$ is derived in the paper for the case of an actin filament; it has units $(\mathrm{m\,mol^{-1}})$ .
